# Calpain-2 inhibition or deletion enhances levels of the transcription Factor, MEIS2, and stimulates neurogenesis

**DOI:** 10.1101/2025.09.22.677817

**Authors:** Michel Baudry, Tristan Reece, Roxana Shahi, Katelyn Ta, Xiaoning Bi

## Abstract

Neurogenesis takes place in the subventricular zone (SVZ) and in the dentate gyrus (DG) of the hippocampus of many adult mammalian species. Recent findings indicate that calpain-2 could participate in neurogenesis regulation through the truncation of the transcription factor, Myeloid Ecotropic Viral Integration Site 2 (MEIS2). The present study aimed to test the effects of calpain-2 inhibition/deletion on MEIS2 levels and neurogenesis in adult mice. Two-to-three month-old mice were injected with a selective calpain-2 inhibitor, NA-184, and sacrificed 24 h later. In addition, two-to-three month-old conditional calpain-2 knock-out (C2KO) and calpain-1 knock-out (C1KO) mice were used. Levels of MEIS2 and of cell markers for neurogenesis were analyzed using immunohistochemistry and western blots. Dendritic spines in hippocampal neurons were also analyzed by Golgi staining. Acute treatment of wild-type (WT) mice with NA-184 increased levels of MEIS2 in various brain structures. It also increased numbers of neurons immunopositive for Ki67 and DCX, two markers for neurogenesis, in both the SVZ and DG. MEIS2 levels were elevated in C2KO mouse brain, while they were decreased in C1KO mouse brain. Compared to those in WT mice, neurons from C2KO mice exhibited a decrease in the number of filipodia spines and an increase in the number of mushroom spines, while those from C1KO exhibited opposite changes. These findings further emphasize the critical and opposite roles of calpain-1 and calpain-2 in brain functions in general, and in neurogenesis in particular with MEIS2 as a major downstream mediator. These findings also underline previous conclusions that calpain-1 promotes spine maturation and synaptic plasticity while calpain-2 hinders spine maturation and synaptic plasticity. These results indicate that calpain-2 inhibition/deletion results in increased neurogenesis, as well as in increased maturation of dendritic spines, potentially due to increased levels and activation of MEIS2.

**Graphical Abstract:** 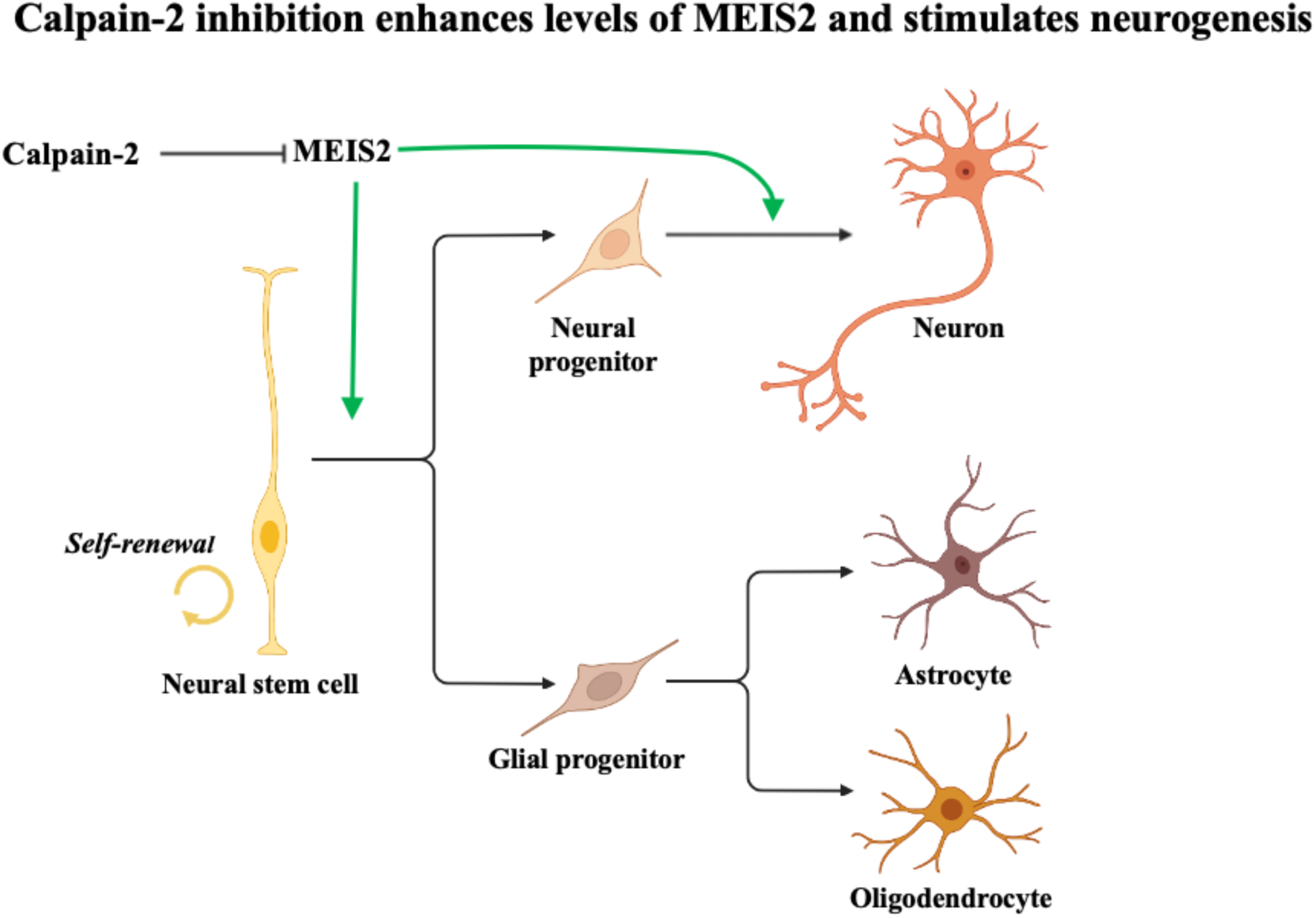

## Introduction

It is now well accepted that, in mammals, neurogenesis persists throughout the lifetime [1], although the rate of proliferation declines in adulthood and further decreases with aging [2, 3]. Adult neurogenesis is mostly observed in two regions of the adult mammalian brain, the subventricular zone (SVZ) and the dentate gyrus (DG) of the hippocampus [4, 5]. In these two regions, endogenous stem cells give rise to neural progenitor cells (NPC), which then proliferate, differentiate and migrate [2]. Some of these cells differentiate into neurons, which integrate into existing neural circuits in the hippocampus and olfactory bulbs [6, 7].

A number of studies have indicated that calpains, a family of calcium-dependent proteases, play an important role in neurogenesis, cell proliferation, survival, migration and differentiation, through their cleavage of proteins involved in these processes [8–10]. In the brain, the most abundant calpain isoforms are calpain-1 and calpain-2, which differ in their calcium sensitivity and their PDZ interacting sequences in their C-terminal domains [11]. We previously demonstrated that calpain-1 played a critical role in neuronal survival, as the number of apoptotic neurons are increased in the brain of calpain-1 knock-out (C1KO) mice during the early postnatal period [12]. Calpain-1 levels are elevated in immature neural progenitor cells, while levels of calpain-2 increase with differentiation, suggesting that calpain-2 activity is required for differentiation [13]. We also observed reduced proliferation of newly generated cells in the DG of C1KO mice [10]. Opposite roles of calpain-1 and calpain-2 in cell adhesion, cell death and cell proliferation/differentiation have been reported in mammary tissue [14]. The role of calpain-2 in neurogenesis has been more difficult to establish due to the lack of calpain-2 knock-out (C2KO) mice, as global calpain-2 deletion is embryonically lethal [15], and until recently, of selective calpain-2 inhibitors.

A recent study reported that the transcription factor, myeloid ecotropic viral integration site 2 (MEIS2) is a target for calpain-2 [16]. MEIS2 is involved in almost all aspects of CNS development, including NPC proliferation, neuronal maturation and synaptogenesis [17–23]. These results raised the possibility that calpain-2 could regulate neurogenesis through the truncation of MEIS2. Our laboratory has produced a series of selective calpain-2 inhibitors, which we have shown to be neuroprotective in a number of acute models of neuronal injuries [24–27]. The present study tested the effects of acute administration of one of these selective calpain-2 inhibitors, NA-184, on brain levels of MEIS2 and neurogenesis in adult mice. As we have also generated conditional calpain-2 knock-out (C2KO) mice with deletion of calpain-2 in excitatory neurons of the forebrain, we compared MEIS2 levels and spine maturation in wild-type (WT), C1KO and C2KO mice. The results indicate that calpain-2 deletion or inhibition results in increased brain levels of MEIS2, increased neurogenesis and increased spine maturation in the adult mouse brain.

## Materials and Methods

### Animals

Animal experiments were conducted in accordance with the principles and procedures of the National Institutes of Health Guide for the Care and Use of Laboratory Animals. All protocols were approved by the local Institutional Animal Care and Use Committee of Western University of Health Sciences. This study also adheres to internationally accepted standards for animal research, following the 3Rs principle. The ARRIVE guidelines were employed for reporting experiments involving live animals, promoting ethical research practices. We used C57Bl/6 (WT), CamKII-Cre^+/-^ CAPN2^loxP/loxP^ (C2CKO), calpain-1 knock-out (C1KO) mice (generous gift from Dr. Athar Chisthi; Tufts University, Boston). C2CKO mice were generated by crossing male Cre^-/-^CAPN2^loxP/loxP^ with female Cre^+/-^ CAPN2^loxP/loxP^ (these mice were purchased from the Riken Institute, Japan). All mice used in this study were males, on a C57Bl/6 background, about 3 months of age and weighing about 30 g. They were kept in a 12 h day/night cycle and were monitored by the WesternU vivarium personnel as well as with the laboratory personnel.

### NA-184

NA-184 ((S)-2-(3-benzylureido)-N-((R,S)-1-((3-chloro-2-methoxybenzyl)amino)-1,2-dioxopentan-3-yl)-4-methylpentanamide) was synthesized by Element (Santa Clara, CA). NA-184 (1 mg/kg in 5% DMSO) was injected intraperitoneally (i.p.) twice at time 0 and 8 h later. Animals were sacrificed 24 h after the initial injection. Investigators performing the analysis of the various outcomes were blind to the treatment (NA-184 or vehicle). Structure and properties of NA-184 have recently been reported [27].

### Immunohistochemistry (IHC)

In cases where whole-brain sections were used, mice were deeply anesthetized after NA-184 administration and perfused intracardially with freshly prepared 4% paraformaldehyde in 0.1 M phosphate buffered saline (pH 7.4). After perfusion, brains were removed and immersed in 4% paraformaldehyde at 4°C for post-fixation, then in 15% and 30% sucrose at 4°C for cryoprotection. Coronal frozen sections of each brain were prepared at 20 μm thickness using a cryomicrotome (CM 1950, LEICA). In cases where half-brain sections were used, mice were deeply anesthetized after NA-184 administration and sacrificed. Brains were removed and halved; one hemisphere was immediately placed into freshly prepared 4% paraformaldehyde in 0.1 M phosphate buffered saline (pH 7.4) for 48 h of post-fixation before being processed in the same manner described above. The other hemisphere was used for western blot analyses.

Free-floating tissue sections were rinsed twice in 0.1 M phosphate buffered saline (PBS) for 15 min, then once in 0.1M PBS with 0.3% Triton-X100 (0.3% PBST) for 30 min. Sections were blocked for 1 h at room temperature (RT) in 5% goat serum (GS) in 0.3% PBST [28]. After blocking, free-floating tissue sections were incubated in primary antibody (MEIS2 (Proteintech), Doublecortin (DCX) (Santa Cruz), Ki67 (Invitrogen) and Olig2 (Invitrogen)) diluted in a solution of 1% bovine serum albumin (1% BSA) with 0.3% PBST overnight at 4 °C. The following day, sections were briefly washed 3 times for 5 min each at RT in PBS followed by incubation with secondary antibodies (AlexaFluor goat anti-rabbit 594, AlexaFluor goat anti-mouse 488 (Invitrogen)) diluted in a solution of 1% BSA with 0.3% PBST for 2 h at RT. Sections were then briefly washed 4 times for 5 min each at RT in PBS and mounted using Vectashield mounting media with DAPI (H-1200, Vector Laboratories). Brain sections were imaged using tile scans on a Zeiss LSM 880 confocal laser-scanning microscope. Identically sized regions of interest (ROI’s) were outlined on each image using the “draw rectangle” function of ZenBlue that captured the entirety of the brain area being analyzed, and mean fluorescence intensity (MFI) was measured. Four sections were stained and analyzed for each brain, and the results were averaged.

### Western Blots

Mice were deeply anesthetized with isoflurane 24 h after NA-184 administration and sacrificed. Brains were removed and dissected to isolate areas of interest, then immediately frozen on dry ice and stored at −80 °C until further processing. Whole homogenates were produced by homogenizing tissue in RIPA buffer. Protein concentrations were determined with a BCA protein assay kit (Pierce).

Proteins were separated by SDS-PAGE followed by transfer to PVDF membranes (Cat# IPFL00010, Millipore) [28]. After incubation with 3% BSA/TBS (Tris-buffered saline) at RT for 1 h, membranes were incubated with primary antibodies overnight at 4 °C (MEIS2 (Proteintech), ß-actin (Sigma-Aldrich)). Following incubation in primary antibodies, membranes were washed with TBST (Tris-buffered saline with Tween-20) 3 times for 10 min each, at RT, and then incubated with secondary antibodies (IRDye secondary antibodies, mouse (LI-COR Biosciences) for 2 h at RT with gentle shaking. Membranes were then washed with TBST (Tris-buffered saline with Tween-20) 4 times for 10 min each, before a final wash in TBS at RT. The specific protein bands were analyzed with the Odyssey imaging system and the LI-COR Image Studio Software.

### Dendritic spine analysis

Golgi impregnation was performed according to the protocol outlined in the FD Rapid GolgiStain Kit (FD Neurotechnologies). One hundred micrometer sagittal sections of the forebrain were cut and mounted. Images of dendritic branches of hippocampal CA1 pyramidal neurons were acquired using a Zeiss light microscope with a 60× objective. The number of spines located on randomly selected dendritic branches was counted manually by an investigator blind to genotype. Spine density was calculated by dividing the number of spines on a segment by the length of the segment and was expressed as the number of spines per micrometer of dendrite. Five to seven distal dendritic branches between 10 and 20 μm in length were analyzed and averaged for a slice mean. The following spine parameters were measured: length, head diameter and neck width. Spines were classified as i) mushroom: large mushroom-shaped with head diameter > 0.6 µm, neck/head ration < 0.5 and ii) filopodia spines: thin, no clear head, elongated shape (head diameter < 0.6 µm); stubby: short and thick without a distinct neck; long, thin protrusions (> 2 µm).

## Results

### 1. NA-184 administration increases MEIS2 levels and neurogenesis in both SVZ and DG

We first tested the effects of systemic administration to young adult mice (2-3 months-old) of NA-184 (ip, 1 mg/kg, twice at 8 h interval) on levels of MEIS2 in SVZ and DG after 24 h. We used immunohistochemistry to analyze MEIS2 expression levels in the SVZ and the DG (Fig. 1). Results clearly indicated that MEIS2 levels were increased by about 40% in both areas.

**Figure 1:**
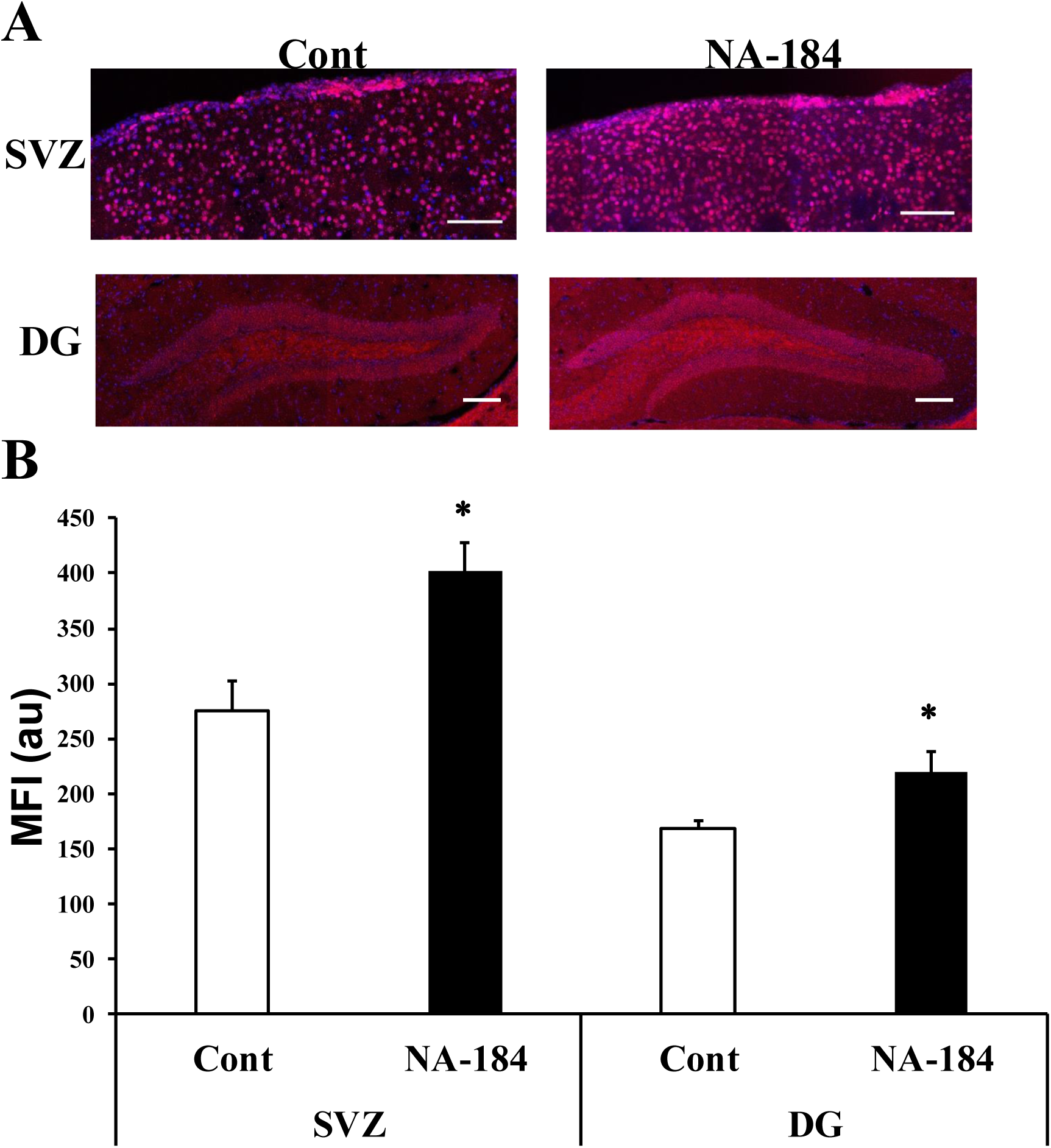
NA-184 treatment increases MEIS2 expression in SVZ and DG. Adult mice were injected (ip) twice (8-h interval) with NA-184 (1.0 mg/kg) or vehicle and sacrificed 24 h later. Brains were processed for immunohistochemistry, using an MEIS2 antibody. Sections were also stained with DAPI (blue). **A.** Representative images of MEIS2 staining in the SVZ and DG. Scale bar = 100 µm. **B.** Quantification of the mean fluorescence intensity (MFI) in the SVZ and DG. Results are means ± S.E.M. of 6 mice. * p <0.01 (Unpaired two-tailed t-test).

We then determined the number of cells immunopositive for Ki67 (Fig. 2) and doublecortin (DCX) (Fig. 3), two commonly used markers for neurogenesis, associated with proliferating cells and immature neurons, respectively, in these 2 brain areas.

**Figure 2:**
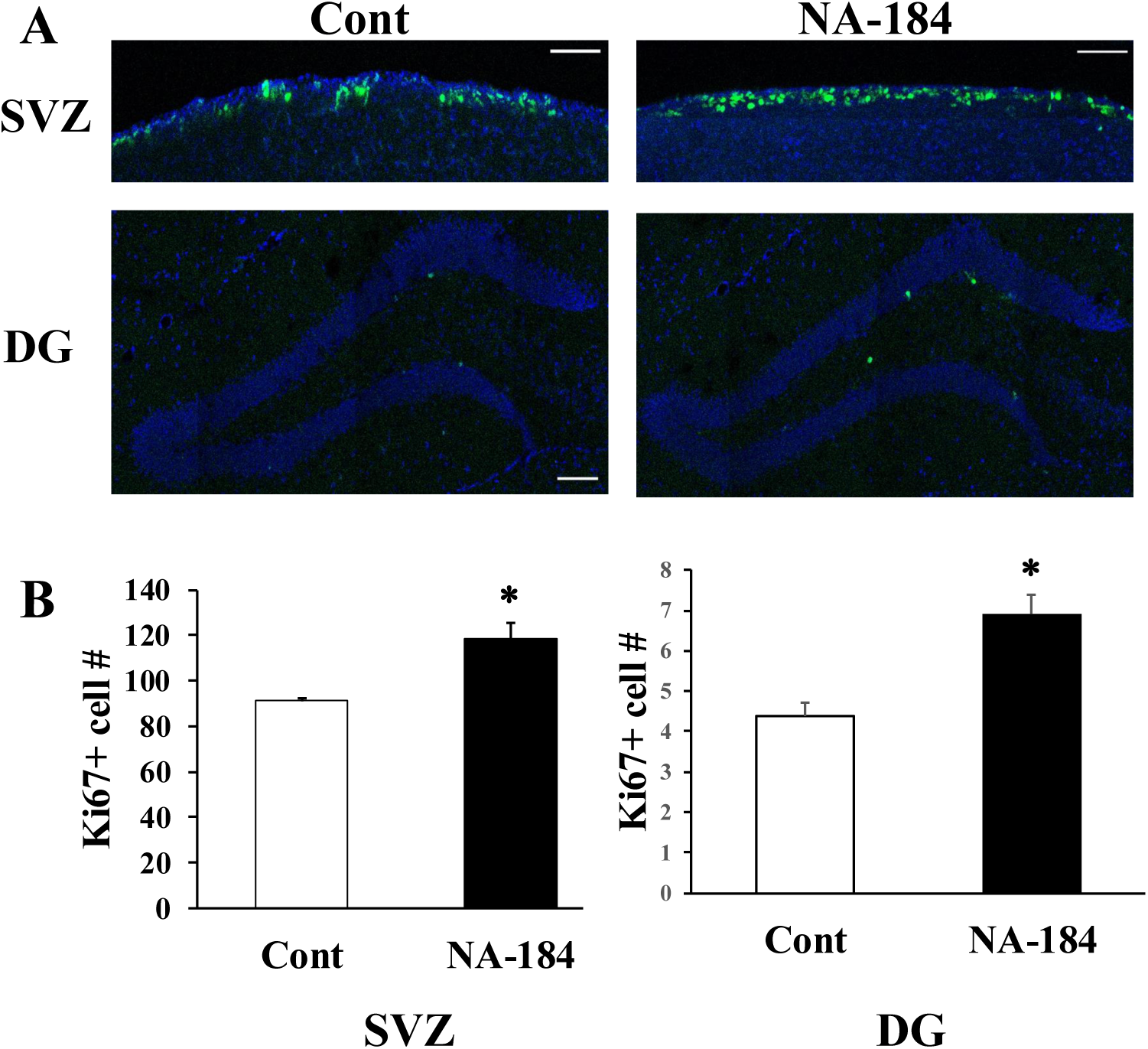
NA-184 injection increases the number of Ki67+ cells in SVZ and DG. Adult mice were injected (ip) twice (8-h interval) with NA-184 (1.0 mg/kg) or vehicle and sacrificed 24 h later. Brains were processed for immunohistochemistry, using an Ki67 antibody. The number of cells labeled with Ki67 were counted in both SVZ and the DG of the hippocampus. **A.** Representative images of Ki67 staining on the SVZ and DG. Scale bar= 100 µm. **B.** Quantification of the number of Ki67+ cells in the SVZ and DG. Results are means ± S.E.M. of 6 mice. * p<0.05, ** p<0.01 (Unpaired two-tailed t-test).

**Figure 3:**
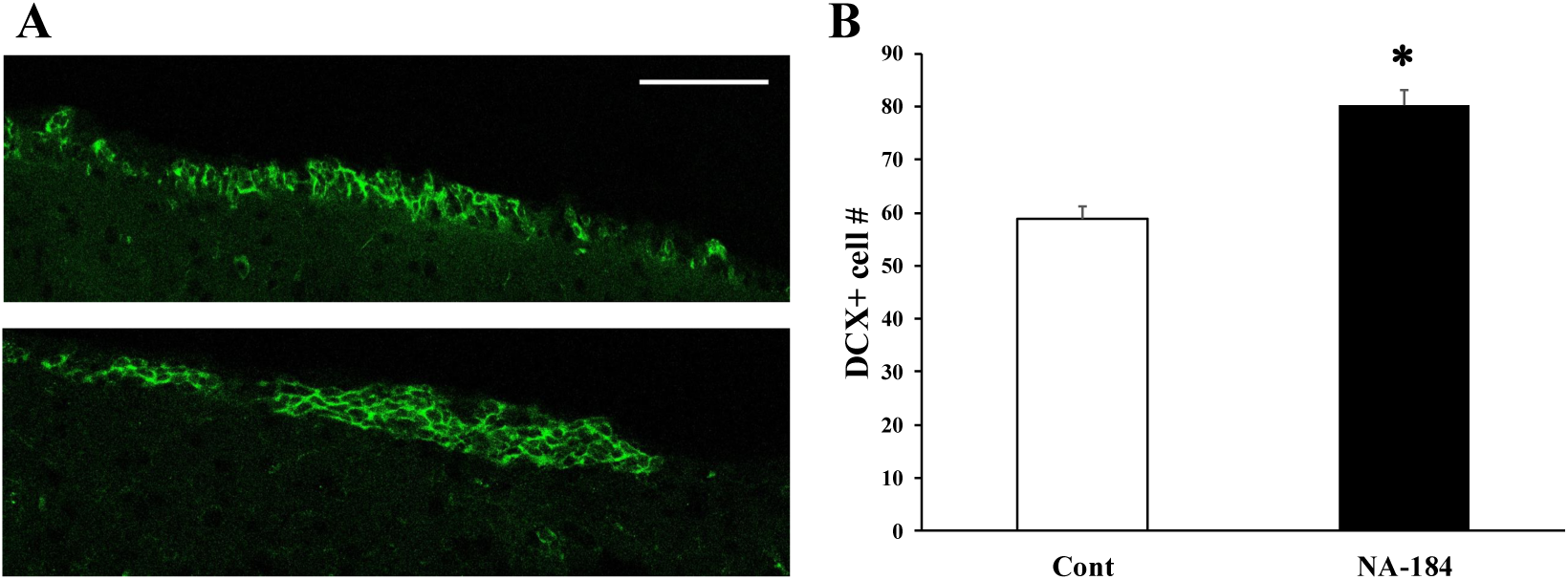
NA-184 injection increases the number of DCX+ cells in SVZ. Adult mice were injected (ip) twice (8-h interval) with NA-184 (1.0 mg/kg) or vehicle and sacrificed 24 h later. Brains were processed for immunohistochemistry, using an antibody against DCX. **A.** Representative images of DCX staining in the SVZ. Scale bar = 100 µm. **B.** Quantification of the number of cells labeled with Ki67 in the SVZ. Results are means ± S.E.M. of 6 mice. * p<0.01 (Unpaired two-tailed t-test).

The number of Ki67-immunopositive cells was increased by about 30-40% in both the SVZ and the DG (Fig. 2). The number of DCX-immunopositive cells was also increased by about 30-40% in the SVZ (Fig. 3). Due to the very low number of DCX-immunopositive cells in the DG, we could not quantify the effects of NA-184 in this structure.

### 2. Changes in MEIS2 levels in mice with deletion of calpain-1 or calpain-2

The availability of mice with deletion of calpain-1 or calpain-2 provided the opportunity to determine the respective roles of these 2 calpain isoforms in the regulation of MEIS2 levels. We used both IHC and western blots to determine the levels of MEIS2 in various brain structures, including CA1, DG and cortex (Figs. 4,5).

**Figure 4:**
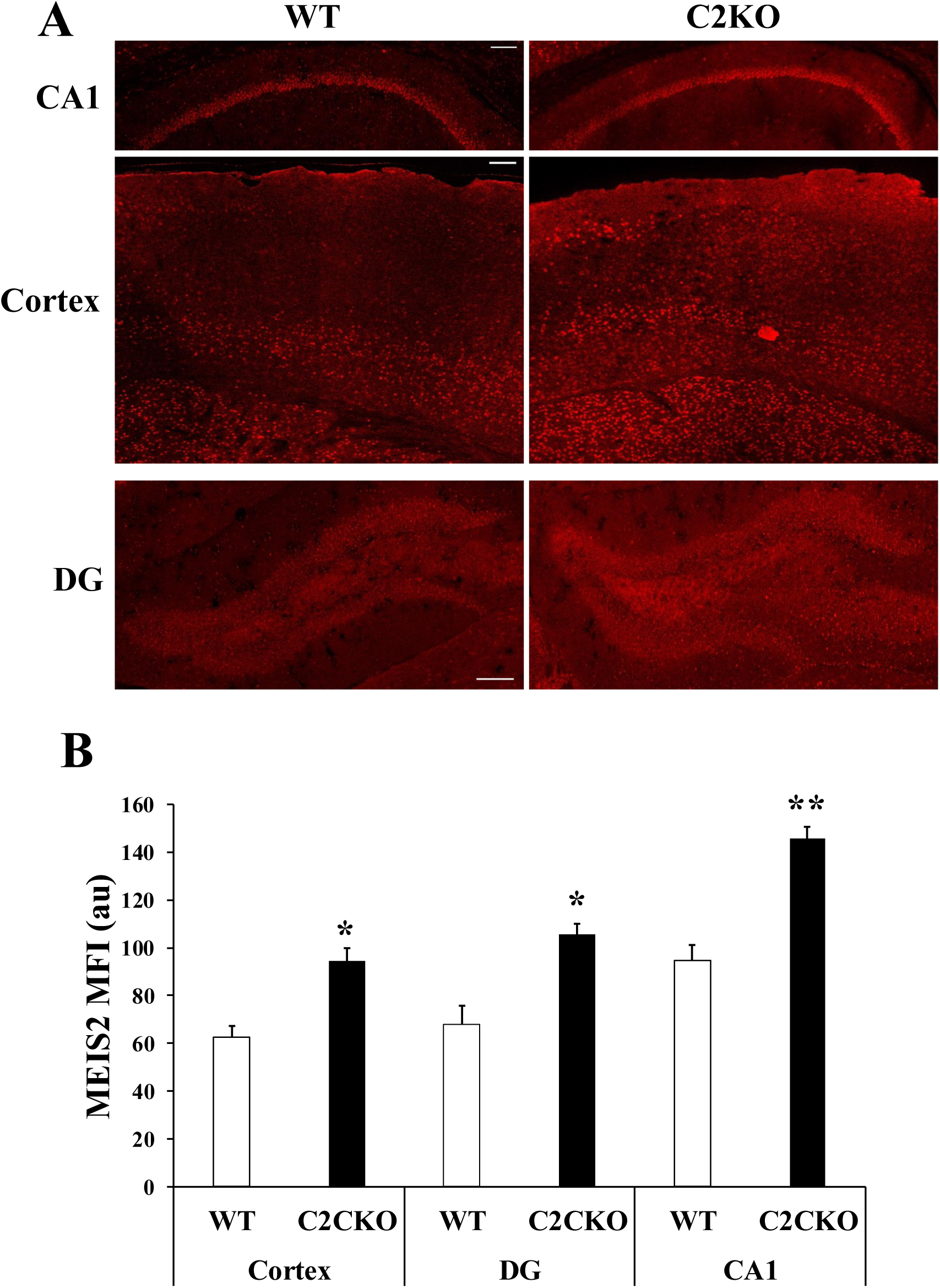
Increased expression of MEIS2 in brain of C2KO mice. Adult WT and C2KO mice were sacrificed, and half the brains were processed for IHC for MEIS2. **A.** Representative images of MEIS2 expression in various brain regions. **B.** Quantification of the changes in MEIS2 expression from images similar to those shown in A. Results are means ± S.E.M. of 4 animals. * p<0.05 and ** p<0.01 (One-way ANOVA followed by unpaired two-tailed t-test).

**Figure 5:**
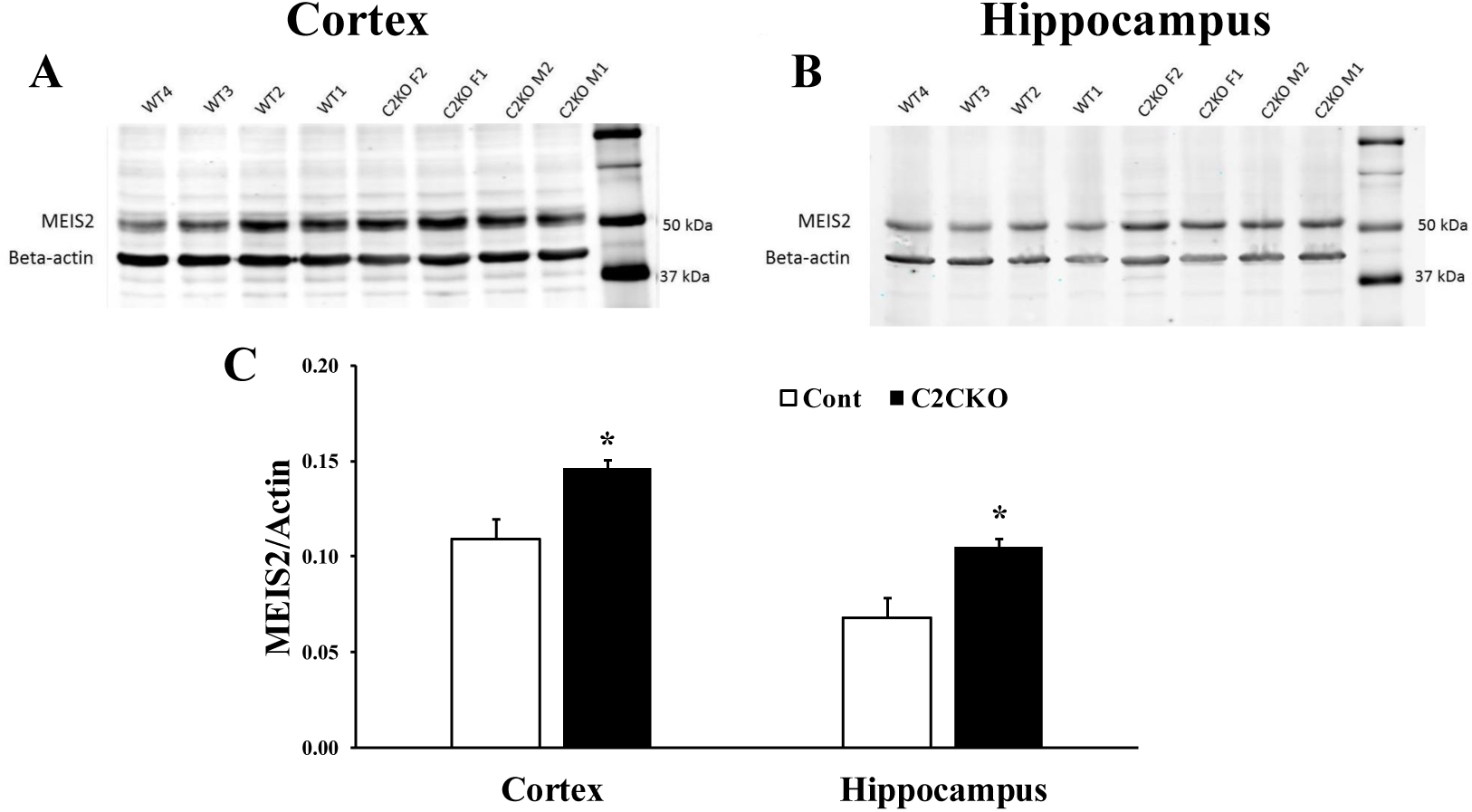
Increased expression of MEIS2 in brain of C2KO mice. Adult WT and C2KO mice were sacrificed, and half the brains were processed for immunoblots for MEIS2. Cortex and hippocampus were dissected and processed for western blots. **A.** Western blots in samples from cortex. **B.** Western blots in samples from hippocampus. **C.** Quantification of the blots. Results are means ± S.E.M. of 4 animals. * p<0.01 (Unpaired two-tailed t-test).

Because calpain-2 deletion is selective for the excitatory neurons of the forebrain, it was not possible to analyze the subventricular zone in these mice. MEIS2 levels were elevated in all brain regions analyzed both with IHC or western blots, and the increase was about 30-40%, a magnitude similar to what was found with acute injection of NA-184, thus confirming that the effects of NA-184 are due to calpain-2 inhibition.

To verify that MEIS2 was a selective calpain-2 substrate, we determined the changes in MEIS2 levels in hippocampus and cortex of C1KO mice (Fig. 6). Interestingly, MEIS levels were not increased but were significantly reduced in both hippocampus and cortex by about 30%.

**Figure 6:**
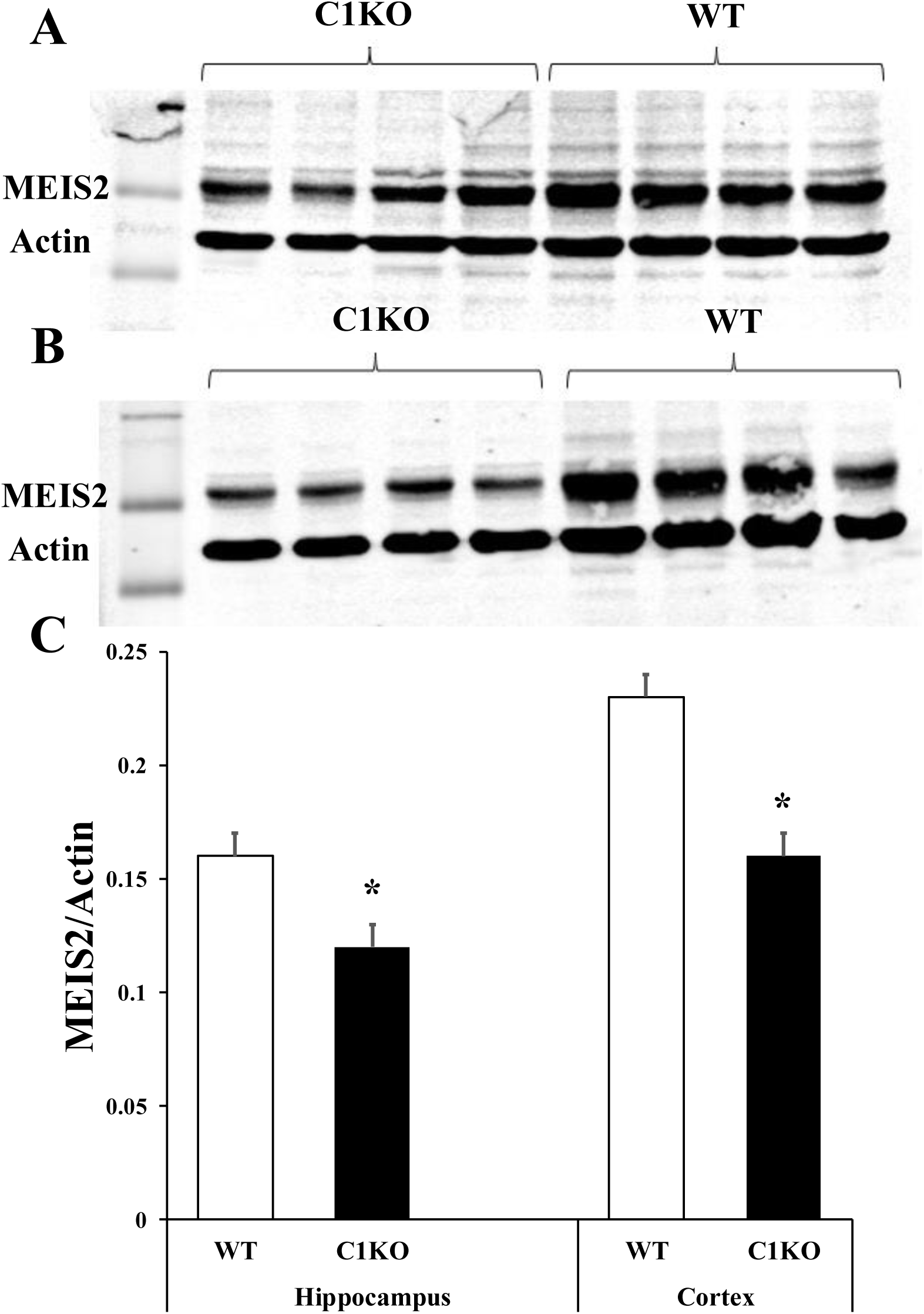
Decreased levels of MEIS2 in adult C1KO brain. Adult WT and C2KO mice were sacrificed, and cortex and hippocampus were dissected and processed for immunoblot for MEIS2. **A.** Western blots in samples from the hippocampus. **B.** Western blots in samples from the cortex. **C.** Quantification of the western blots. Results are means ± S.E.M. of 4 animals. * p<0.05 (Unpaired two-tailed t-test).

### 3. Changes in dendritic spines in C2KO and C1KO mice

As MEIS2 is involved in neuronal maturation, we analyzed potential changes in dendritic spine morphology in both C2KO and C1KO hippocampal neurons. Dendritic spines were classified as filopodia or mushroom and their numbers were determined both in basal and apical dendrites of pyramidal neurons of field CA1 (Fig. 7).

**Figure 7:**
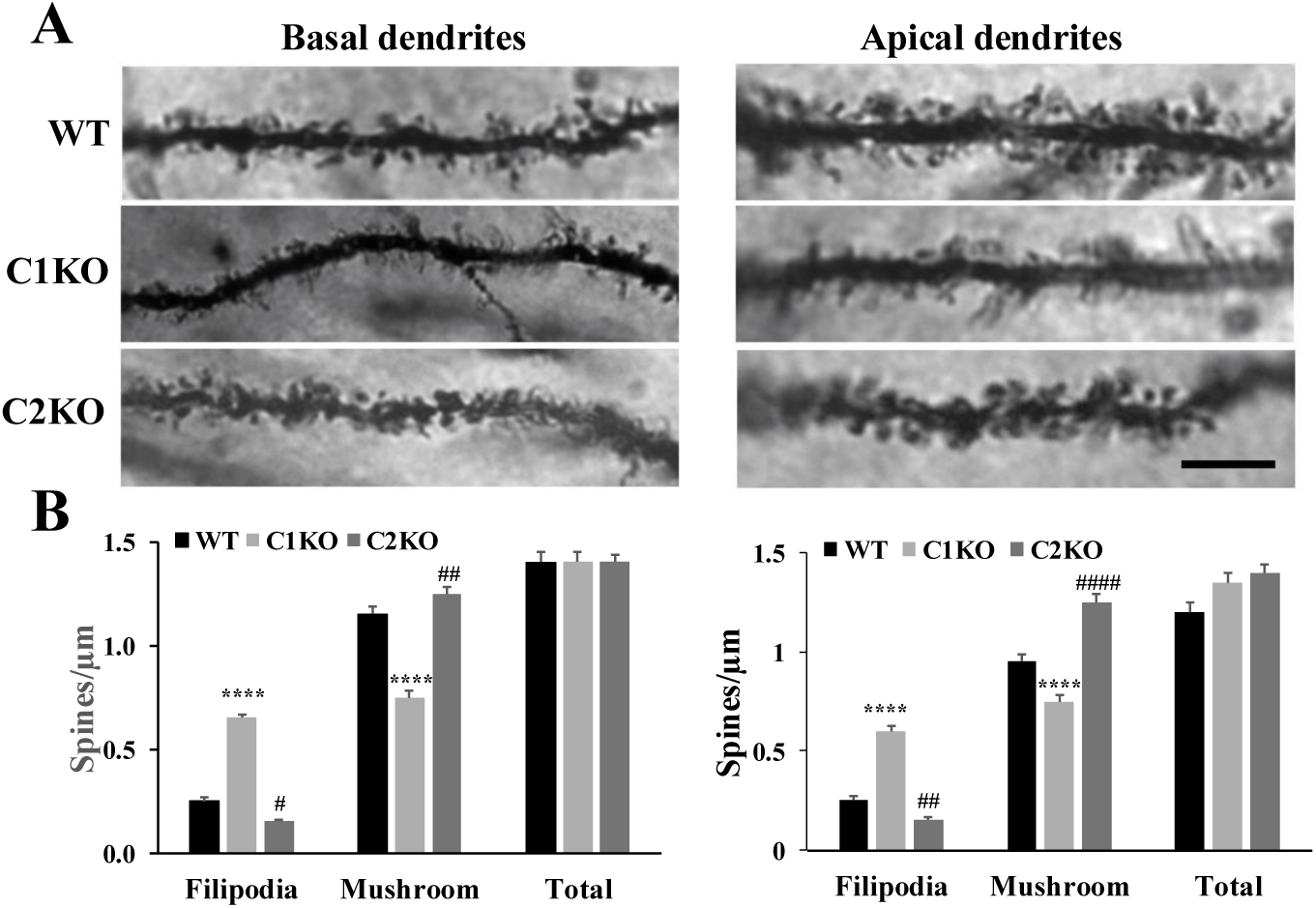
Dendritic spine analysis in basal and apical dendrites of field CA1 in adult C2KO and C1KO mice. Golgi staining was performed in adult brains from WT, C1KO and C2KO mice. Dendritic spines were classified as filopodia or mushrooms and their number quantified in apical and basal dendrites of CA1. **A.** Representative images of the Golgi staining in basal and apical dendrites of a CA1 pyramidal neuron. Scale bar = 10 µm. **B.** Quantification of the number of spines in basal and apical dendrites. Results are means ± S.E.M. of 4 mice per group. * p<0.05, **** p<0.001, as compared to WT. # p<0.05, ## p<0.01, #### p<0.001, as compared to both WT and C1KO (ANOVA followed by unpaired t-test).

In both C1KO and C2KO mice, the total numbers of spines were similar to those found in WT mice. However, the distribution of spines between filopodia and mushroom was changed in opposite directions for C1KO and C2KO mice, as compared to WT mice. Thus, there was an increase in filopodia and a decrease in mushroom spines in both basal and apical dendrites of field CA1 in C1KO mice, as compared to WT mice. On the other hand, there was a decrease in filopodia and an increase in mushroom spines in C2KO mice.

## Discussion

Our results confirm and expand the previous findings that the transcription factor MEIS2 is a target for calpain-2 [16]. Calpain-2 activity was shown to decrease in NPCs during differentiation, resulting in enhanced MEIS2 levels [16]. Both acute injection of a selective calpain-2 inhibitor, NA-184, or deletion of calpain-2 in excitatory neurons of the forebrain were associated with enhanced MEIS2 levels in various neuronal populations, including the hippocampus and cortex. Acute injections of NA-184 enhanced MEIS2 levels in both the SVZ and the DG of the hippocampus, two structures known to exhibit continuous neurogenesis in adult mice. As expected from MEIS2 functions, elevated MEIS2 levels in these two structures were associated with enhanced neurogenesis, as reflected by increased numbers of Ki67 and DCX-immunopositive cells in the SVZ and the Ki67-immunopositive cells in the DG of the hippocampus.

While MEIS2 levels were elevated in brains of C2KO mice, as expected, they were surprisingly decreased in brains from C1KO mice. First, these results confirm the specific role of calpain-2 in the regulation of MEIS2 levels through truncation. We previously showed that calpain-2 levels were not affected in brains of C1KO mice [9], which eliminates the possibility that the decrease in MEIS2 levels in these mice could be due to an increase in calpain-2 expression. We also identified a number of genes regulated by calpain-1 in the brain [29]. In particular, Hspa1, which codes for the chaperone protein, HSP70, was markedly downregulated in brains from C1KO mice. As HSP70 could regulate the correct folding of MEIS2, it is conceivable that decreased expression of HSP70 could result in decreased levels of MEIS2 in C1KO mice. Second, they further expand the range of functions in which calpain-1 and calpain-2 play opposite roles [30], as calpain-1 appears to be pro-neurogenesis while calpain-2 could decrease neurogenesis. These results corroborate our previous findings of decreased proliferation of newly generated neurons in the DG of adult C1KO mice [10].

MEIS2 has also been proposed to play a significant role in neuronal maturation, as knock-down of MEIS2 in prefrontal cortex results in cognitive deficits [31] [17]. Our results indicating that dendritic spines are more mature in hippocampal neurons from C2KO mice support a role for MEIS2 in dendritic spine maturation. Furthermore, the opposite changes observed in C1KO mice also fit well with such a role for MEIS2 in spine maturation. Several mechanisms could account for the observed effects. MEIS2 regulates the expression of a number of genes, including protein kinases such as Cdk5, which have been shown to participate in the regulation of spine morphology [32]. We previously showed that calpain-2 targets PTEN [33], a phosphatase also implicated in synaptogenesis [34]. Independent of the molecular mechanisms underlying these effects of calpain-1 and calpain-2 on spine morphology, it remains clear that the balance between calpain-1 and calpain-2 activity plays a critical role in the maturation and maintenance of spine morphology.

The current results have important implications for the potential beneficial effects of selective calpain-2 inhibitors for the treatment of numerous disorders associated with impaired neurogenesis and synaptogenesis. We have already shown that selective calpain-2 inhibitors have beneficial effects for the treatment of acute glaucoma [24], severe traumatic brain injury (TBI) and repeated mild concussions [35], as well as seizure-induced brain inflammation and cognitive impairment [26]. In addition, a selective calpain-2 inhibitor was found to enhance synaptic plasticity and learning and memory in young adult mice [36]. It is possible that the beneficial effects we observed under all these conditions could, at least in part, be related to the effects of the inhibitor on neurogenesis and synaptogenesis. In addition, it is also possible that this novel function of calpain-2 in the regulation of neurogenesis could support the use of calpain-2 inhibitor for the treatment of disorders associated with reduced neurogenesis, including age-related neurodegenerative disorders, such as Alzheimer’s disease, Parkinson’s disease and Amyotrophic Lateral Sclerosis (ALS). Importantly, a very recent study confirmed the presence of proliferating neural progenitors in the adult human hippocampus, suggesting the possibility of using NA-184 to stimulate neurogenesis in adult and aged humans [37.]

## Conclusion

The results of the present study clearly demonstrate that MEIS2 is a target of calpain-2 but not of calpain-1. They also demonstrate that mice treatment with a selective calpain-2 inhibitor increases MEIS2 levels in brains leading to increase in neurogenesis. As MEIS2 is also involved in synaptogenesis, calpain-2 inhibition or deletion could result in increased dendritic spine maturation. The results have significant implications for the use of selective calpain-2 inhibitors for the treatment of a number of neurodegenerative disorders.

## Limitations

While our study clearly shows that calpain-2 inhibition stimulates neurogenesis and spine enlargement through MEIS 2 enhancement, there could be parallel pathways contributing to this effect. The potential functional outcomes of this effect were not evaluated in our study, although they are consistent with the cognitive enhancement we previously reported for selective calpain-2 inhibition. The long-term effects of calpain-2 inhibition could disturb the balance between proliferation versus differentiation. The effects on other cell types, including oligodendrocytes remain to be evaluated.

## Acknowledgments

The authors wish to acknowledge the contributions of Drs. Yubin Wang and Jiandong Sun, who performed the spine analysis reported in Figure 6.

## Author contributions

Tristan Reece, Roxana Shahi and Katelyn Ta performed the experiments. Xiaoning Bi and Michel Baudry organized the experiments, generated the funding for the studies and wrote the manuscript.

## Animal rights

As indicated above, animal experiments were conducted in accordance with the principles and procedures of the National Institutes of Health Guide for the Care and Use of Laboratory Animals. All protocols were approved by the local Institutional Animal Care and Use Committee of Western University of Health Sciences.

## List of abbreviations

AD: Alzheimer’s disease
ALS: amyotrophic sclerosis
BSA: bovine serum albumin
C1KO: calpain-1 knock-out
C2KO: calpain-2 knock-out
DCX: doublecortin
DG: dentate gyrus
MEIS2: myeloid ecotropic viral integration site 2
MFI: mean fluorescence intensity
NPC: neural progenitor cells
NA-184: (S)-2-(3-benzylureido)-N-((R,S)-1-((3-chloro-2-methoxybenzyl)amino)-1,2-dioxopentan-3-yl)-4-methylpentanamide
PBS: phosphate buffered saline
PD: Parkinson’s disease
TBI: traumatic brain injury
SVZ: subventricular zone

